# Uncovering protein-protein interactions of the human sodium channel Nav1.7

**DOI:** 10.1101/2023.04.18.537407

**Authors:** Xuelong Zhou, Jing Zhao

**Author notes:** Correspondence: Jing Zhao, Molecular Nociception Group, Wolfson Institute for Biomedical Research, University College London, London WC1E 6BT, UK. or Xuelong Zhou, Department of anesthesiology and perioperative medicine, The First Affiliated Hospital of Nanjing Medical University, Nanjing, Jiangsu 210029, China.

## Abstract

The voltage-gated sodium channel Nav1.7 plays a crucial role in the initiation and propagation of pain signals. Our previous study has successfully identified the interacting proteins of mouse Nav1.7 (mNav1.7). In this study, we aimed to further elucidate the protein-protein interactions associated with human Nav1.7 (hNav1.7). Stable epitope (TAP)-tagged HEK293 cells expressing hNaV1.7 were utilized for the identification of hNav1.7-interacting proteins. The hNaV1.7-associated complexes were isolated through tandem affinity purification and further characterized by mass spectrometry.

Bioinformatics analysis was carried out using the PANTHER classification system. Electrophysiological recording was performed to assess Nav1.7 current. Tap-tagged hNav1.7 was expressed effectively in HEK293 cells, exhibiting normal functional Nav1.7 currents. A total of 261 proteins were identified as interactors of hNav1.7, mainly located across the cell membrane and cytoplasm, and primarily involved in biological processes related to protein translation and expression. Comparison between human and mouse Nav1.7-interacting proteins revealed shared proteins (such as Eef1a1, Eef2, Tcp1, Cct2, Cct3, Cct5, Cct6a, and Cct7) as well as protein families (such as kinesin and Rab GTPases family). Knockdown of two of the shared interacting proteins, CCT5 and TMED10, resulted in reduced Nav1.7 current density. In conclusion, the protein interactions of hNaV1.7 were successfully mapped in the current work. These novel findings offer essential insights into the regulatory mechanisms that govern Nav1.7 function.

**Significance statement:** Chronic pain affects approximately 20% of the world’s population and is a global major public health problem. Nav1.7 has been recognized as a promising target for novel analgesics. However, the drug development process for Nav1.7 is challenging. A thorough understanding of the regulatory mechanism of Nav1.7 would greatly assist in the development of its analgesic drugs. Our previous work successfully mapped the mNav1.7 protein interactions. In the current study, the interacting proteins of hNav1.7 were further defined. Our findings provide important implications for the development of Nav1.7-based analgesics for human use.

## Introduction

In recent decades, the prevalence of individuals suffering from pain has significantly increased due to the aging population and other social and psychological factors. According to statistical data, the number of individuals experiencing chronic pain has surpassed the total number of individuals suffering from heart disease, cancer, and diabetes (Steglitz et al., 2012). Despite the utilization of various pain management techniques, such as physiotherapy, medications, and surgery, pain control remains inadequate for approximately 40% of patients (Johannes et al., 2010). Moreover, many medications have severe side effects, including the risk of addiction, respiratory depression, and other complications. As a result, developing novel analgesics and treatments with minimal side effects has emerged as a crucial focus of pain research.

To date, nearly ten mutant genes have been discovered to be implicated in the generation of aberrant human pain phenotypes. For example, SCN9a∼SCN11a, which encode sodium voltage-gated channel alpha subunit Nav1.7∼Nav1.9 (Sexton et al., 2018), and PRDM12, which encodes the PR domain zinc finger protein 12 (Chen et al., 2015). Among these genes, only SCN9a presents a sufficient and necessary condition for the changes in pain phenotype. Gain-of-function mutations in the SCN9a gene have been found in patients with hyperalgesia, such as primary erythema limb pain (Yang et al., 2004), paroxysmal severe pain (Fertleman et al., 2006), and small fiber neuropathy (Faber et al., 2012). Conversely, loss-of-function mutations in the SCN9a gene were identified in patients with congenital insensitivity to pain. Importantly, individuals with SCN9a gene mutations have been found to exhibit normal motor, cardiovascular, and cognitive functions (Cox et al., 2006). In preclinical studies, the expression of Nav1.7 was found to be increased in primary sensory neurons of inflammatory (Liang et al., 2013), neuropathic (Sadamasu et al., 2014), and postoperative pain rodents (Sun et al., 2018).

Furthermore, conditional deletion of Nav1.7 has been shown to induce deficiencies in acute and inflammatory pain (Nassar et al., 2004). These findings indicate that Nav1.7 is indispensable for the generation and transmission of pain signals, making it a promising target for the development of new analgesic agents.

Over the past few decades, a significant amount of effort has been dedicated to the development of selective Nav1.7 blockers. However, the outcomes have been less than satisfactory. Some blockers are ineffective due to their inability to penetrate the blood-brain barrier, while others exhibit minimal analgesic efficacy but significant adverse effects (Yekkirala et al., 2017). Recent structural biology research has revealed that human voltage-gated sodium channel subtypes exhibit remarkable structural similarity (Huang et al., 2017). Developing selective Nav1.7 blockers directly poses a significant challenge. An alternative approach, involving the indirect manipulation of Nav1.7 biological processes such as protein synthesis or intracellular trafficking, resulting in the reduction of its membrane expression, could be effective. Previously, we employed tandem affinity purification (TAP) and liquid chromatography-tandem mass spectrometry (LC-MS/MS) to characterize the protein interactions of mouse Nav1.7 (mNav1.7) (Kanellopoulos et al., 2018). In the present study, we utilized similar approaches to map the protein-protein interactions of human Nav1.7 (hNav1.7). Our findings provided crucial insights into the biological mechanism of Nav1.7 by comparing the interacting proteins of hNav1.7 to those of mNav1.7.

## Materials and methods

### Extraction of gene expression data from BIOGPS

The gene expression data pertaining to Nav1.7 biological process in human dorsal root ganglion (DRG) and HEK293 cells was obtained from BIOGPS (http://biogps.org/#goto=welcome). Specifically, the expression data for key genes associated with this biological process in human DRG was derived from the Geneatlas U133A-gcrma database, while the data for these core genes in HEK293 cells was extracted from the NCI60 on U133A-gcrma database. To quantify the relative gene expression level, the gene expression levels were normalized against the internal reference gene GAPDH.

### Generation of a stable TAP-tagged hNaV1.7-expressed HEK293 cell line

A stable HEK293 cell line expressing hNaV1.7 with a TAP tag (TAP-hNaV1.7) was generated according to the previous description (Koenig et al., 2015). The TAP tag consists of a peptide sequence (SRK DHL IHN VHK EEH AHA HNK IEN LYF QGE LPT AAD YKD HDG DYK DHD IDY KDD DDK) that is inserted immediately upstream of the stop codon in the SCN9A expression construct. The TAP tag contains a HAT (histidine affinity tag) domain and three FLAG tags, which can be detected using anti-HAT and anti-FLAG antibodies. Immunohistochemistry, western blotting, and electrophysiological analyses were conducted to evaluate the expression and function of TAP-tagged hNaV1.7.

### Immunocytochemistry

Poly-D-lysine coated coverslips were placed in 24-well plates and HEK293 cells were incubated for 24 hours. The cells were subsequently permeabilized using cold methanol for 10 minutes, followed by fixation in cold acetone for 1 minute. The fixed cells were treated with a blocking buffer composed of 0.3% Triton X-100 and 10% goat serum in PBS and left to incubate at room temperature for 30 minutes. After washing with PBS, the cells were exposed to an Anti-FLAG antibody (1:500, F1804, Sigma) overnight at 4°C. The secondary antibody (Alexa Fluor 488, 1:2000, A11017, Invitrogen) was subsequently added and incubated with the washed cells for 2 hours at room temperature. After washing with PBS, the coverslips were mounted using a DAPI-containing mounting solution and visualized under a fluorescence microscope.

### Western blotting

Harvested cells were homogenized in lysis buffer consisting of Tris 20 mM, NaCl 100 mM, 1% DDM, 0.2% CHS, protease inhibitor cocktail, and adjusted to pH 7.4. The lysate was centrifuged at 20,000g for 10 minutes at 4°C to collect the supernatant. The protein concentrations in the supernatant were determined using the Pierce BCA protein assay kit, and a total of 40 μg protein was separated by SDS-PAGE gel. The separated proteins were then transferred onto a polyvinylidene difluoride membrane. The membranes were blocked with 5% nonfat milk for 30 minutes before being treated with primary antibodies, including anti-FLAG (1:1000, Sigma, F1804), anti-Nav1.7 (1:500, Origene, TA329033), anti-CCT5 (1:1000, Origene, TA308298), and anti-TMED10 (1:1000, Origene, TA306375), overnight at 4°C. Following TBS-Tween washes, the membranes were incubated with the secondary antibody (goat anti-mouse or goat anti-rabbit IgG-HRP, 1:2000; Jackson Immuno-Research Laboratories) for 2 hours at room temperature. The protein bands on the membrane were visualized using a chemiluminescence reagent system.

### Small interfering RNA

Small interfering RNAs (siRNAs) were designed to target the human CCT5 and TMED10 genes, based on their genomic sequences (NM_012073 for CCT5 and NM_006827 for TMED10). The sequences of CCT5 siRNAs were: 5-CCGAGUCCAUUGUUAAUGATT-3 and 5-UCAUUAACAAUGGACUCGGTT-3. The sequences of TMED10 siRNAs were: 5-GGCGAUGUGACUAUAACAATT-3 and 5-UUGUUAUAGUCACAUCGCCTT-3. A scrambled sequence was also designed as a mismatch control. The siRNAs were transfected into HEK293 cells using Lipofectamine 2000 (Invitrogen). The effect of knockdown was validated by western blot.

### Electrophysiology and patch-clamp recordings

Whole-cell patch-clamp recording was conducted according to previously established protocols (Emery et al., 2015). an Axopatch 200B amplifier and a Digidata 1322a digitizer (Axon Instruments) were utilized with Clampex software (version 10, Molecular Devices) for operation. Filamented borosilicate microelectrodes (GC150TF-10, Harvard Apparatus) were pulled with a micropipette puller (Model P-97, Sutter Instruments) and fire-polished to achieve resistances between 2.5 and 4 MOhm.

The pipette solution consisted of NaCl 10 mM, CSF 140 mm, EGTA 1.1 mm, MgCl2 1 mM, HEPES 10 mm, and pH 7.3, while the extracellular solution featured NaCl 140 mM, MgCl2 1 mM, KCl 3 mM, CaCl2 1 mM, HEPES 10 mm, and pH 7.3. Following immersion of the electrode in the extracellular solution, the pipette offset potential was zeroed. The electrode was slowly approached to the cell membrane, and negative pressure was applied to achieve a giga-seal. Electrode capacitance was compensated, and gentle suction was used to break the membrane, allowing for whole-cell recording to be established. Cells were held at -100 MV for 2 minutes prior to initiating the experiment protocol.

Normalized peak currents at various applied voltage steps were measured to generate the current density-voltage curve. The voltage-dependent activation and inactivation data were fit using the following Boltzmann equations: y=(A2+(A1-A2)/(1+exp((Vh-x)/k)))*(x-Vrev) for activation and y=(A1-A2)/(1+exp((x-Vh/k))+A2 for inactivation. The parameters A1, Vh, x, Vrev, and k are the maximal amplitude, potential of half-maximal activation, clamped membrane potential, reversal potential, and a constant, respectively.

### Tandem affinity purification

The collected cells were homogenized in the lysis buffer. The lysate was then centrifuged at 20,0000 g for 30 minutes at 4°C, and the resulting supernatants were collected and incubated overnight at 4°C with Anti-FLAG resin (Sigma, A2220). The Anti-FLAG resin was subsequently collected and washed, after which the interesting proteins were competitively eluted using 3XFLAG peptide (Sigma, F4799). The eluent was then incubated overnight at 4°C with Ni-NTA beads (Qiagen, 36111), which were subsequently collected, washed, and eluted using imidazole buffer. The eluent was further concentrated and purified using size exclusion chromatography (SEC). A portion of the eluent was preserved to confirm the protein mass spectrometry (MS) results.

### SEC and MS

The eluent containing the target proteins underwent filtration using a Centricon filter with a 100 kD cutoff (EMD Millipore). The Centricon filter was then centrifuged at 1,200g for 10 minutes until the sample volume was reduced to less than 1 mL. The concentrated sample was subsequently loaded onto the sample loader of the Superose 6 chromatography column (Cytiva) using a syringe. The AKTA chromatography system facilitated the separation of the sample within the column. The isolated proteins were subjected to protein MS analysis.

### Analysis of hNav1.7-interacting proteins

The Panther classification system was utilized to analyze the interacting proteins of hNav1.7. Data pertaining to mNav1.7-interacting proteins were obtained from our previous work (Kanellopoulos et al., 2018) (https://www.embopress.org/action/downloadSupplement?doi=10.15252%2Fembj.201796692&file=embj201796692-sup-0002-TableEV1.Xlsx).

### Data analysis

The data were presented as the mean ± standard deviation (SD). The comparison between two groups was analyzed using an unpaired Student’s t-test. Multiple group comparisons were assessed using repeated-measures analysis of variance (ANOVA). A statistically significant difference was considered when *P* < 0.05. The electrophysiological data were extracted using pCLAMP software, and the statistical analysis was performed using Origin 9.0 software.

## Results

### Genes related to Nav1.7 are similarly expressed in human DRG neurons and HEK293 cells

Nav1.7 is primarily expressed in primary sensory neurons, such as DRGs. However, obtaining sufficient human DRG tissues to map Nav1.7 protein interactions is challenging. HEK293 cells are commonly used as a tool to overexpress hNav1.7 and possess essential genes involved in protein synthesis, intracellular trafficking, and membrane expression of Nav1.7. To further validate this, we compared the expression levels of critical genes involved in Nav1.7 protein folding, intracellular trafficking, membrane anchoring, and post-transcriptional modification between HEK293 cells and human DRGs. The results showed that HEK293 cells and human DRG neurons possess similar gene expression profiles (Figure 1A-1F) (Bao, 2015).

**Figure 1.**
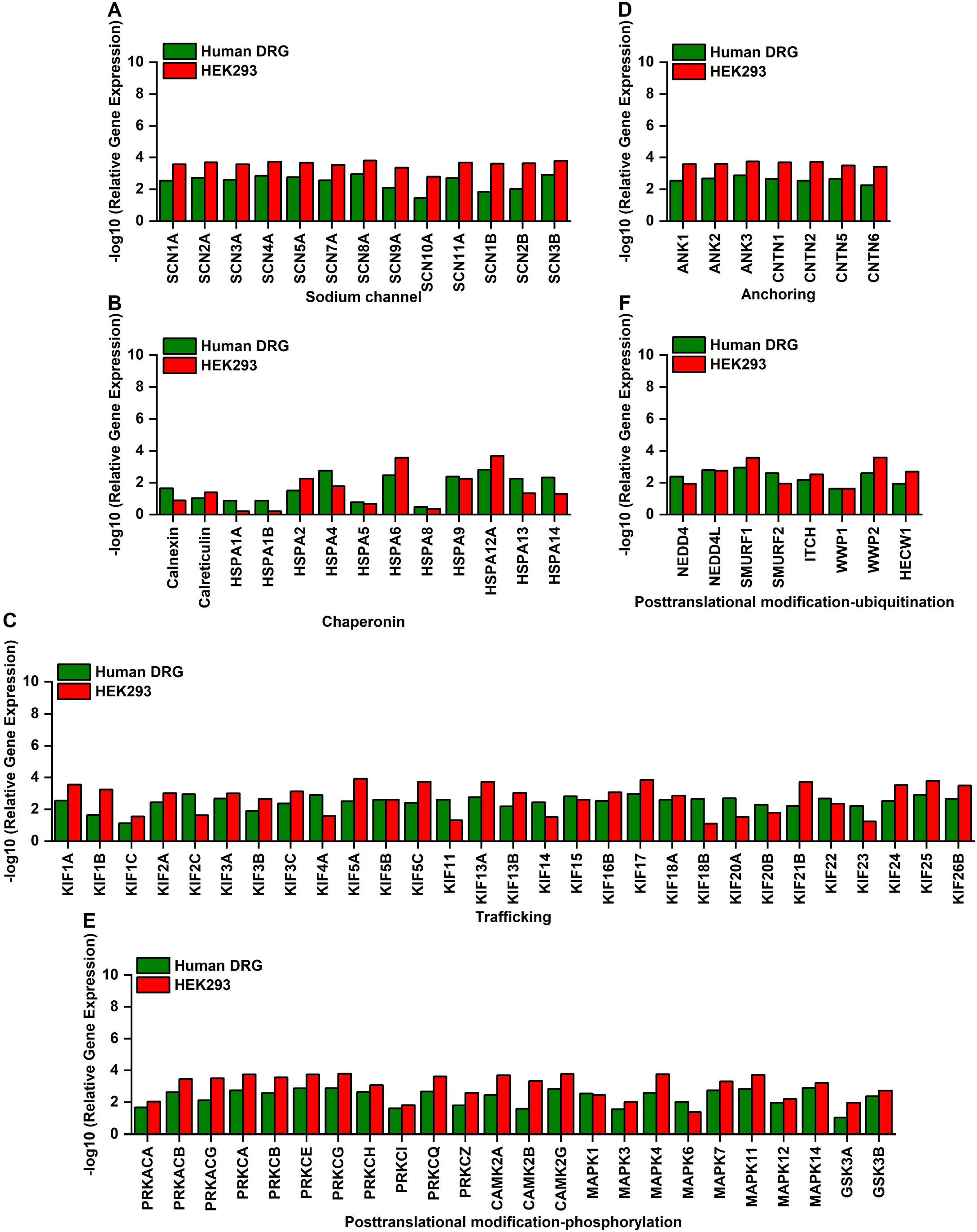
Comparison of Nav1.7-related gene between human DRG and HEK293 cells. **A**. The gene expression levels of sodium channel alpha subunits and beta subunits in human DRG and HEK293 cells. **B**. The gene expression levels of Nav1.7 related-chaperonins in human DRG and HEK293 cells. **C**. The gene expression levels of kinesins in human DRG and HEK293 cells. **D**. The gene expression levels of Nav1.7 related-membrane anchoring proteins (ankyrins and contactins) in human DRG and HEK293 cells. **E and F**. The gene expression levels of Nav1.7-related post-transcriptional modification (phosphorylation, E and ubiquitination, F) proteins in human DRG and HEK293 cells.

### Characterization of hNav1.7 in HEK293 cells

To map out the protein-protein interactions of hNav1.7, a stable TAP-tagged hNav1.7 expressed HEK293 cells was employed (Koenig et al., 2015). The TAP tags have a molecular weight of 5 kD and consist of a poly-HAT and a 3×Flag tag in tandem (Figure 2A). Immunofluorescence staining and western blot revealed that the TAP-tagged hNav1.7 was successfully expressed on HEK293 cells (Figure 2B and 2C). Furthermore, electrophysiological analyses showed that the TAP-tagged hNav1.7 HEK293 cells displayed normal functional currents (Figure 2D-2I).

**Figure 2.**
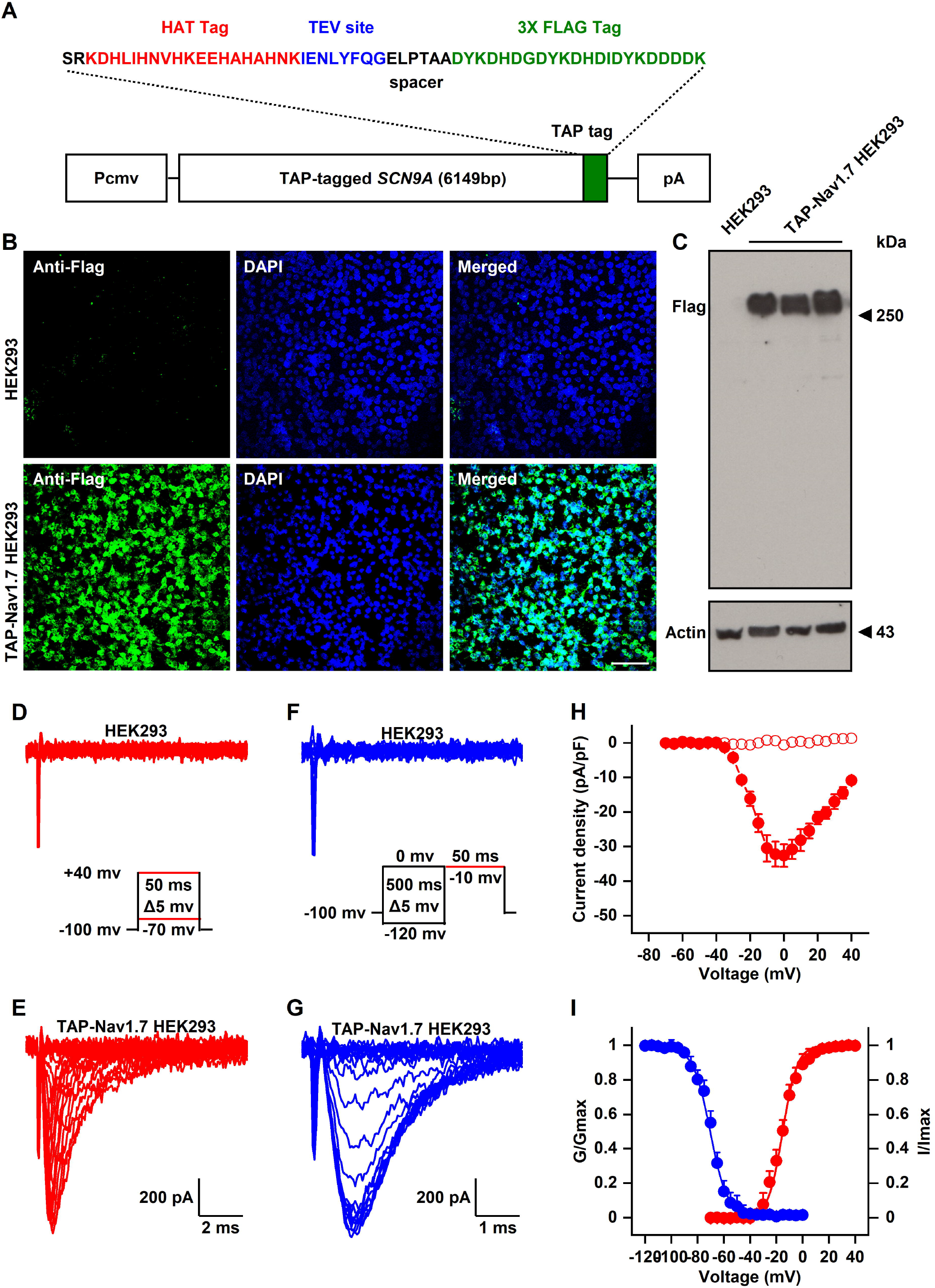
Characterization of TAP-tagged hNaV1.7 in HEK293 cells. **A**. Schematic diagram of TAP-tagged SCN9A cDNA construct. **B**. Immunohistochemistry with an anti-FLAG antibody was conducted on HEK293 cells stably expressing TAP-tagged hNav1.7 and parental HEK293 cells. Scale bar = 100 μm. **C**. Western-blot analysis for TAP-tagged hNav1.7 was performed in HEK293 cells stably expressing TAP-tagged hNav1.7 and parental HEK293 cells. **D and E**. Representative raw current traces of NaV1.7 from HEK293 cells stably expressing the TAP-tagged NaV1.7 and parental HEK293 cells in response to the activation pulse protocol shown. **F and G**. Representative raw current traces of NaV1.7 from HEK293 cells stably expressing the TAP-tagged NaV1.7 and parental HEK293 cells in response to the inactivation pulse protocol shown. **H**. The current-voltage relationship of the channel in HEK293 cells stably expressing the TAP-tagged hNav1.7 and parental HEK293 cells. **I**. The activation and inactivation voltage dependence of the channel in HEK293 cells stably expressing the TAP-tagged hNav1.7.

### Purification of hNav1.7-associated complexes

The TAP-tagged Nav1.7 and its associated protein complex was solubilized in a lysis buffer containing n-dodecyl-D-maltoside (DDM), an ionic detergent that has been shown to purify membrane proteins better than conventional detergents (Sanders et al., 2007). Due to the TAP tag, two consecutive affinity purifications were performed, consisting of a single-step affinity purification (SS-AP) utilizing anti-FLAG resin, followed by a tandem affinity purification using Ni-NTA beads (TAF) (Figure 3A). Western blot results showed that the TAP-tagged hNav1.7 was efficiently precipitated by SS-AP and TAP (Figure 3B). The purified TAP-tagged hNav1.7 and contaminants were further separated by SEC and analyzed via protein mass spectrometry (Figure 3C).

**Figure 3.**
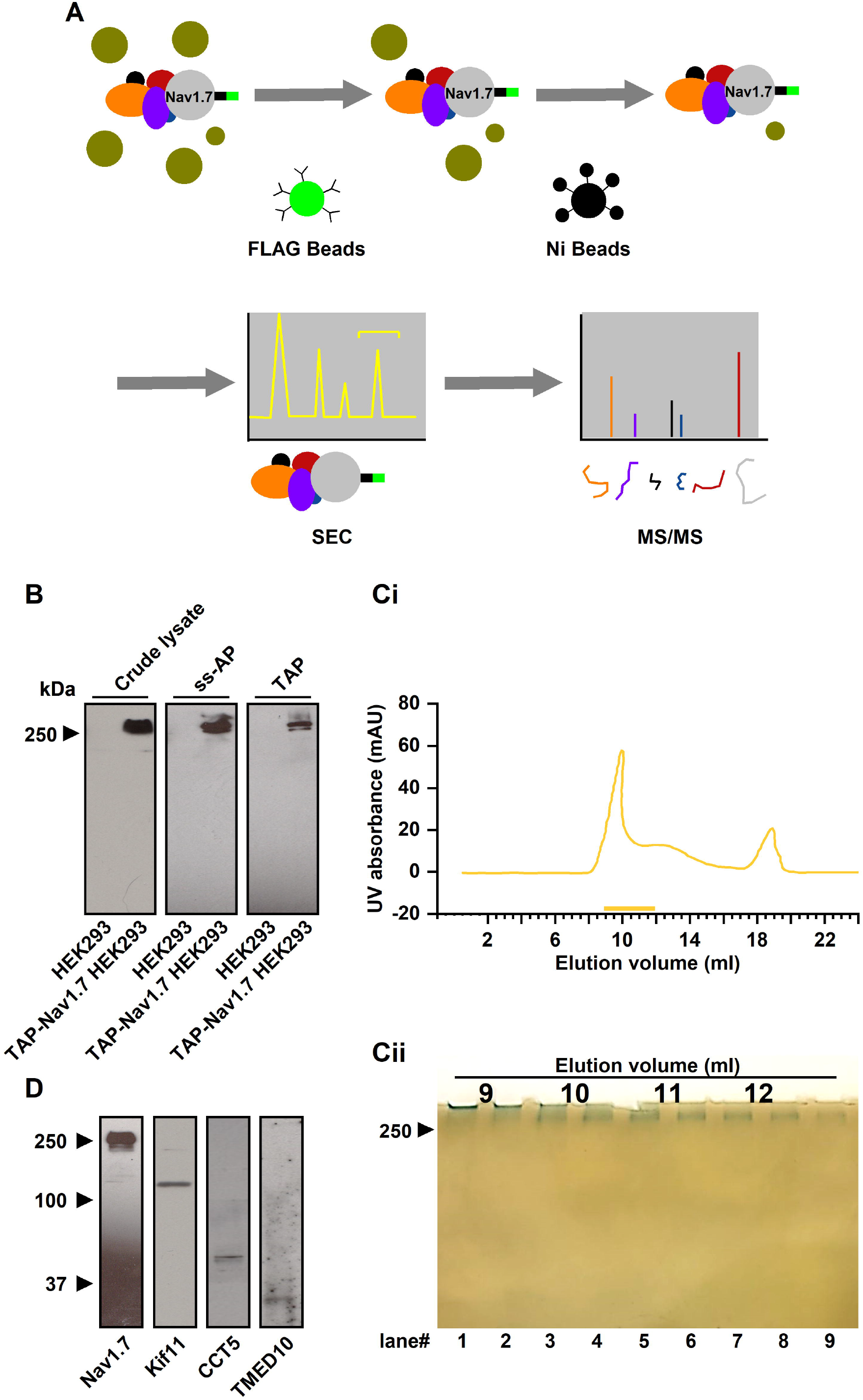
Tandem affinity purification, size exclusion chromatography (SEC), and tandem mass spectrometry (MS/MS) employed for the analysis of TAP-tagged hNav1.7 in HEK293 cells. **A**. Schematic illustrating the procedures involved in the affinity purification, SEC, and MS/MS of TAP-tagged hNav1.7. **B**. TAP-tagged NaV1.7 was detected using western blotting with anti-FLAG antibody after single-step and tandem affinity purification (SS-AP and TAP). **Ci**. A representative size exclusion chromatography of the protein sample. **Cii**. The resolved fractions isolated from size exclusion chromatography visualized using Coomassie blue staining. **D**. The validation of selected Nav1.7 protein interactor candidates Kif11, CCT5, and TMED10, where Nav1.7 complexes were immunoprecipitated with anti-FLAG M2 magnetic beads and then detected with their specific antibodies using Western blotting.

### Identification of hNav1.7-interacting proteins

A total of 261 proteins that physically interact with hNav1.7 were identified through protein mass spectrometry. The top 100 interacting proteins were listed in Table 1. To validate the physical interaction between NaV1.7 and its interacting proteins, three candidates of interest, KIF11, CCT5, and TMED10, were chosen (Figure 3D). PANTHER cell component analysis revealed that these 261 proteins were mainly distributed in the cell membrane and cytoplasm (Figure 4A). Further functional analysis demonstrated that they were primarily involved in protein translation and expression processes, including translation proteins, chaperones, protein modifying enzymes, cytoskeleton proteins, and others (Figure 4B). To compare the differences in Nav1.7 interacting proteins between human and mouse, our previously published mNav1.7 interacting proteins were further investigated. Cell component analysis showed that the mNav1.7 interacting proteins were mainly located in the cell membrane and cytoplasm, with a few protein interactors found on synapses (Figure 4C). Functional analysis showed that the mNav1.7 interacting proteins were engaged in the same processes as the hNav1.7 interacting proteins, which include protein translation and expression processes (Figure 4D).

**Figure 4.**
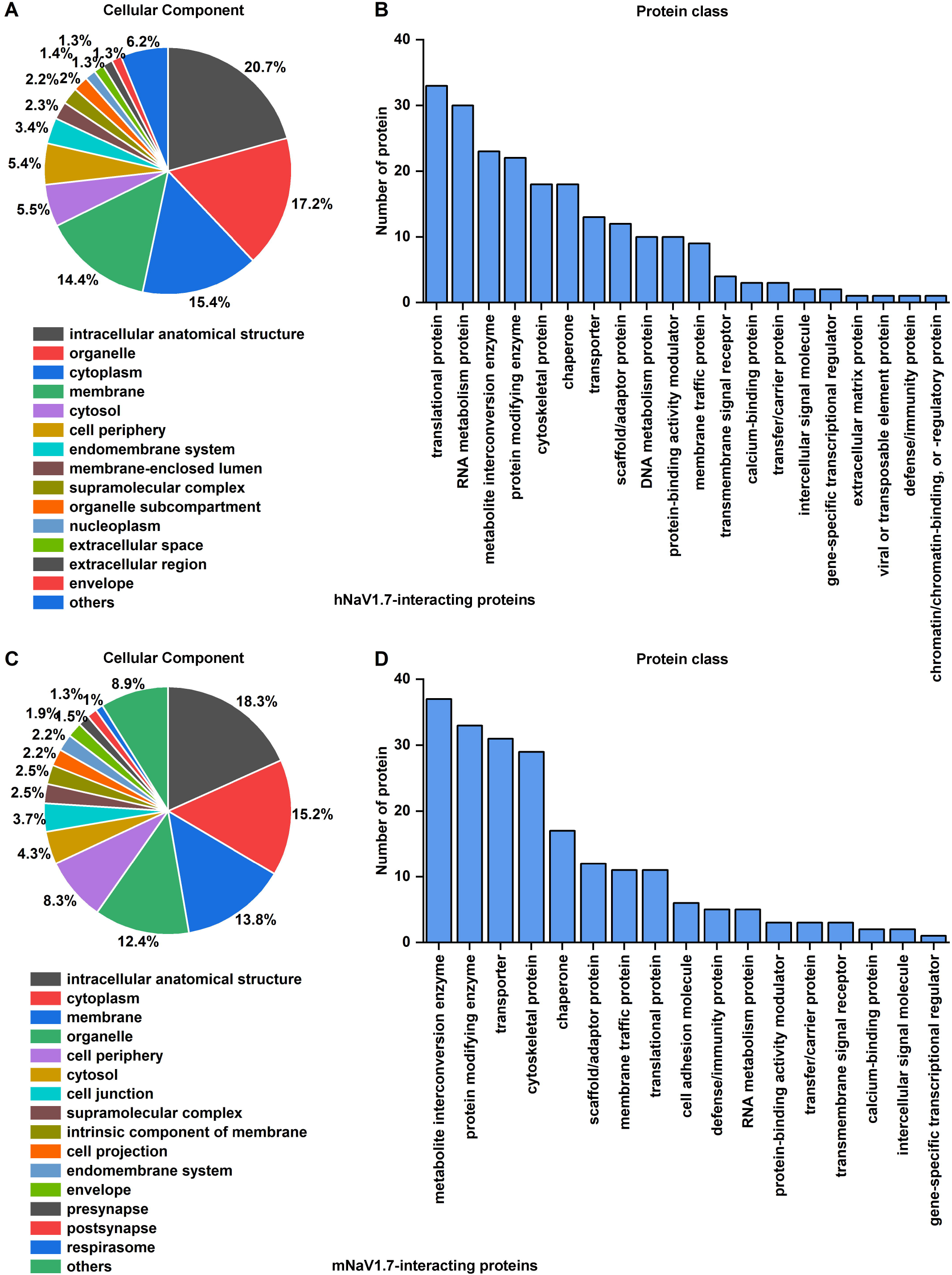
Analysis of NaV1.7 complex proteins using PANTHER Classification System. A. Cellular localization analysis of identified hNav1.7 interacting proteins. B. Protein class analysis of identified hNav1.7 interacting proteins. C. Cellular localization analysis of mNav1.7 interacting proteins. D. Protein class analysis of mNav1.7 interacting proteins.

### Identification of core hNav1.7-interacting proteins

To investigate the similarity between the interacting proteins of hNav1.7 and mNav1.7, a reexamination of their protein interactors in each function category was conducted. The protein interactors of hNav1.7 and mNav1.7 were listed in Table 2 according to their respective function groups. Our analysis revealed that hNav1.7 and mNav1.7 share interacting proteins in various functional groups, such as eukaryotic elongation factors 1a1 and 2 (Eef1a1 and Eef2) in the protein translation group, as well as T-complex protein 1 subunits α, β, γ, ε, ζ, and η (Tcp1, Cct2, Cct3, Cct5, Cct6a, and Cct7) in the chaperone group. Moreover, although the interacting proteins were diverse in some functional groups, they belonged to the same protein family. For instance, hNav1.7 interacting proteins included kinesin family members 5b and 5c (Kif5b and Kif5c) in the cytoskeletal protein group, while mNav1.7 interacting proteins were identified as Kif11. In the protein-binding activity modulator group, Ras-related protein Rab5a was found to interact with hNav1.7, whereas Rab1a was identified as the mNav1.7 protein interactor. As these proteins and protein families demonstrated interactions with both hNav1.7 and mNav1.7, their dependence on neither tissue nor species suggests their potential criticality in regulating Nav1.7.

### Functional verification of core Nav1.7-interacting proteins

To examine whether the core interacting proteins identified above have an impact on Nav1.7 function, two proteins CCT5 and TMED10, which have been previously shown to interact with Nav1.7, were selected for further investigations. The knockdown of CCT5 and TMED10 (∼50%) using siRNA (Figure 5A and 5B) led to a decrease in Nav1.7 current density by approximately 50% and 20% (Figure 5C-5G), respectively, indicating a regulatory role of the core interacting proteins CCT5 and TMED10 in Nav1.7. These Nav1.7 core interacting proteins identified through comparison of hNav1.7 and mNav1.7 interacting proteins may provide insights into the regulatory mechanisms of Nav1.7.

**Figure 5.**
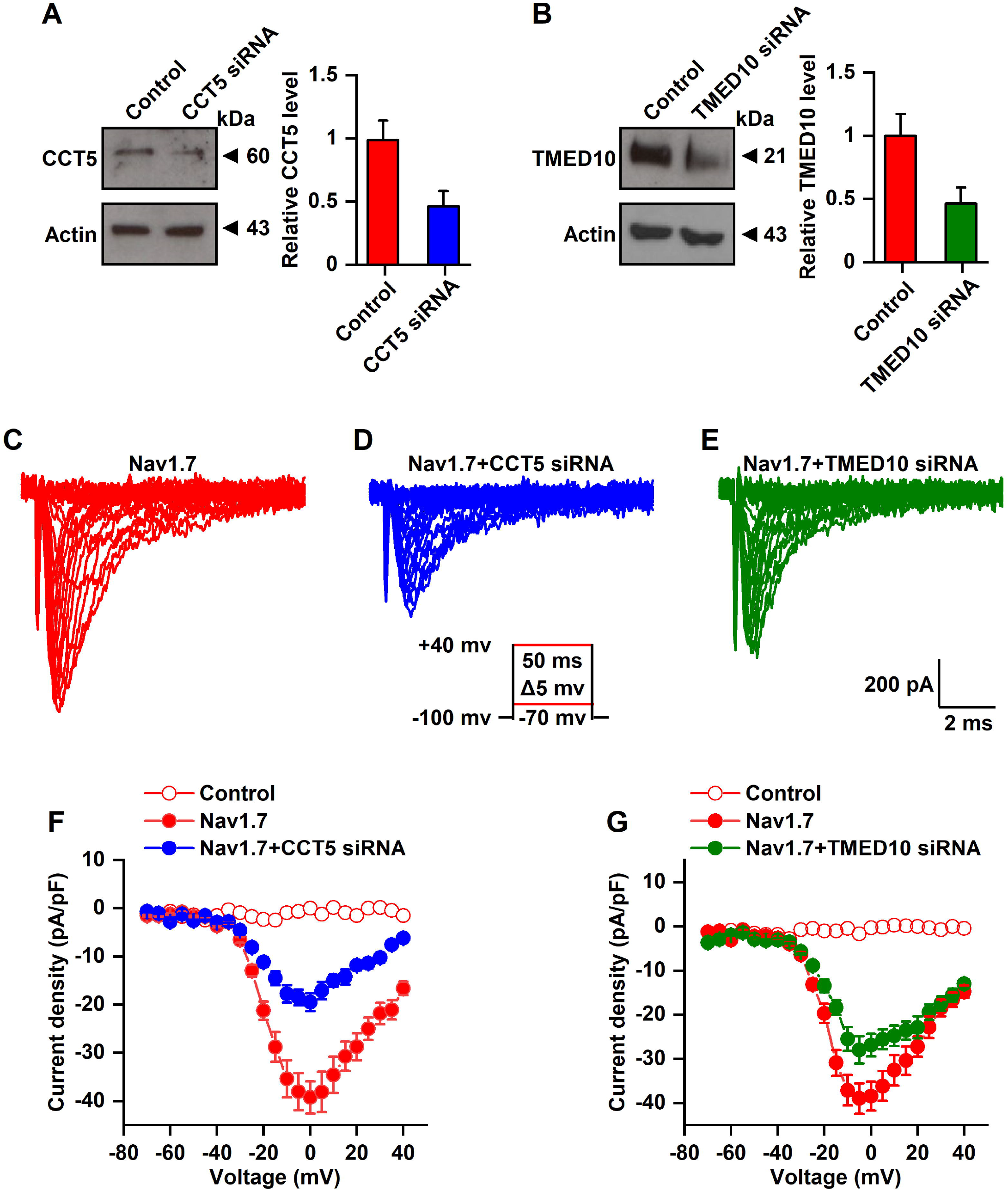
Electrophysiological characterization of NaV1.7 in HEK293 cells following transfection with CCT5 siRNA or TMED10 siRNA. **A**. Expression of CCT5 in HEK293 cells in response to the transfection of CCT5 siRNA. n = 4, *P* < 0.05. **B**. Expression of TMED10 in HEK293 cells in response to the transfection of CCT5 siRNA. n = 4, *P* < 0.05. **C-E**. Representative raw current traces of NaV1.7 from HEK293 cells stably expressing the TAP-tagged NaV1.7 in response to the activation pulse protocol shown. Each trace shows a different condition. **F**. IV plot of Nav1.7 current density in the absence and presence of CCT5 siRNA. Compared with NaV1.7 basal currents (n = 12), CCT5 siRNA transfection significantly reduced the sodium channel density (n = 10, P < 0.01). **G**. IV plot of Nav1.7 current density in the absence and presence of TMED10 siRNA. Compared with NaV1.7 basal currents (n = 10), CCT5 siRNA transfection significantly reduced the sodium channel density (n = 10, P < 0.05).

## Discussion

In the current investigation, we conducted a protein-protein interaction mapping analysis for hNav1.7. The main findings are as follows: Firstly, we identified 261 proteins that interact with hNav1.7 in HEK293 cells; Secondly, we observed that certain proteins or protein families are present within both hNav1.7 and mNav1.7 interactomes; Thirdly, we found that these shared proteins or protein families are involved in the regulation of Nav1.7. Our findings offer valuable insights towards unraveling the regulatory mechanism of Nav1.7.

Nav1.7 has been recognized as a promising target for novel analgesics. However, the drug development process for Nav1.7 is challenging. A thorough understanding of the regulatory mechanism of Nav1.7 would greatly assist in the development of its analgesic drugs. In the past two decades, several molecules, including Crmp2 (Dustrude et al., 2013), Nedd4-2 (Laedermann et al., 2013), and Fgf13 (Yang et al., 2017), have been implicated in the regulation of Nav1.7 transport and degradation. However, the protein-protein interaction network remains undefined. Our previous work successfully mapped the mNav1.7 protein interactions (Kanellopoulos et al., 2018). In the current study, the interacting proteins of hNav1.7 were further defined. By using a stable TAP-tagged hNav1.7 expressed HEK293 cells, we found that 261 proteins interacted with hNav1.7, providing important implications for the development of Nav1.7-based analgesics for human use.

In a study conducted by Kanellopoulos et al. (2018), a total of 267 proteins were identified as the interactome of mNav1.7. Similarly, in the present study, a total of 261 protein interactors were identified for hNav1.7. Cell component analyses revealed that the interacting proteins of mNav1.7 and hNav1.7 were primarily located in the cell membrane and cytoplasm. However, mNav1.7 interacting proteins were also found distributed across synapses. Functional analyses demonstrated that both hNav1.7 and mNav1.7 interacting proteins were primarily involved in the biological process of protein translation and expression, while interacting proteins of mNav1.7 were also implicated in active transmembrane transport processes (transporter). These differences in cell component, function, and even interacting proteins between hNav1.7 and mNav1.7 interacting proteins are determined not only by species, but also by tissue sources. Specifically, mNav1.7 protein interactors were derived from mouse neural tissues, while hNav1.7 protein interactors were derived from non-neural HEK293 cells.

Although the interacting proteins of mNav1.7 and hNav1.7 originate from disparate tissues and species, they nonetheless provide insight into the critical partners necessary for Nav1.7’s biological functions. Within the functional category of protein translation, both mNav1.7 and hNav1.7 were found to interact with Eef1 and Eef2. Additionally, serine/threonine kinase 38 (Stk38) and TGF-beta activated kinase 1 (Tab1) were identified as interacting proteins in the protein modification group of both mNav1.7 and hNav1.7. These shared protein interactors are not limited by tissue or species and may therefore function as the regulatory molecules required for Nav1.7’s function. To test this hypothesis, we selected two shared interacting proteins, CCT5 and TMED10. Our results indicate that the knockdown of CCT5 and TMED10 resulted in a reduction of Nav1.7’s current density, suggesting that these shared interacting proteins may indeed impact Nav1.7’s function.

Some of the proteins identified on the list of mNav1.7 and hNav1.7 interacting partners share membership in the same protein family, despite having distinct identities. For instance, Kif11, an hNav1.7 interacting protein in the cytoskeleton protein group, and Kif5b and Kif5c, which are mNav1.7 interacting proteins in the same group, are members of the kinesin family (Hirokawa et al., 2009). Similarly, Rab5a and Rab1a, which are hNav1.7 and mNav1.7 interacting proteins in the protein-binding activity modulator group, respectively, are members of the Rab GTPases family (Raghavan et al., 2022). The expression of these same family proteins may vary across tissues, as suggested by the observation that Kif5c expression is enriched in mouse nerve tissue but present in low abundance in HEK293 cells (Hirokawa et al., 2009). Consequently, Kif11 may serve as an alternative for Nav1.7 trafficking in HEK293 cells. Notably, these interacting protein families are not specific to any given tissue or species, suggesting that they may play an important role in regulating Nav1.7 function. However, the list of mNav1.7 and hNav1.7 interacting proteins does contain completely distinct proteins. One potential explanation for this observation is the possibility of insufficient experimental data, but it is also likely that these differences are due to species- and tissue-specific factors. When using cell lines for in vitro experiments, it is crucial to consider not only the expression differences of proteins of interest, but also the potential involvement of alternative proteins.

In conclution, the present study mapped the protein-protein interactions of hNav1.7, and identified core protein interactors through comparative analysis with mNav1.7. These findings provide valuable insights into the regulation of Nav1.7 biological function.

## Author contributions

X.L.Z. and J.Z. designed experiments, performed experiments, analyzed data and wrote the manuscript.

## Competing Interests

The authors declare no competing interests.

## Notes

### Competing Interest Statement

The authors have declared no competing interest.

