## Supplementary material for "Uncovering protein-protein interactions of the human sodium channel Nav1.7": Table 1

Table 1. Identified hNaV1.7-associated proteins.

| **Protein** | **Gene** | **Protein name** | **Score** | **Mass** | **Num. of matches** | **Num. of significant matches** | **Num. of sequences** | **Num. of significant sequences** | **emPAI** |
| --- | --- | --- | --- | --- | --- | --- | --- | --- | --- |
| KIF11 | KIF11 | Kinesin-like protein KIF11 | 18792 | 119085 | 531 | 531 | 44 | 44 | 7.08 |
| TBB5 | TUBB | Tubulin beta chain | 3927 | 49639 | 111 | 111 | 13 | 13 | 4.12 |
| TBB4B | TUBB4B | Tubulin beta-4B chain | 3416 | 49799 | 101 | 101 | 13 | 13 | 4.59 |
| TBB2A | TUBB2A | Tubulin beta-2A chain | 3216 | 49875 | 99 | 99 | 10 | 10 | 3.2 |
| ANM5 | PRMT5 | Protein arginine N-methyltransferase 5 | 3083 | 72638 | 98 | 98 | 20 | 20 | 3.25 |
| ALBU | ALB | Serum albumin | 2763 | 69321 | 91 | 91 | 16 | 16 | 2.22 |
| TBB4A | TUBB4A | Tubulin beta-4A chain | 2702 | 49554 | 88 | 88 | 11 | 11 | 3.24 |
| CALX | CANX | Calnexin | 2450 | 67526 | 62 | 62 | 15 | 15 | 2.33 |
| TBA1B | TUBA1B | Tubulin alpha-1B chain | 1862 | 50120 | 40 | 40 | 9 | 9 | 1.35 |
| TBA1A | TUBA1A | Tubulin alpha-1A chain | 1740 | 50104 | 37 | 37 | 9 | 9 | 1.36 |
| MEP50 | WDR77 | Methylosome protein 50 | 1662 | 36701 | 38 | 38 | 6 | 6 | 1.17 |
| PIGS | PIGS | GPI transamidase component PIG-S | 1562 | 61617 | 48 | 48 | 12 | 12 | 1.74 |
| K2C1 | KRT1 | Keratin, type II cytoskeletal 1 | 1539 | 65999 | 47 | 47 | 13 | 13 | 1.56 |
| TAB1 | TAB1 | TGF-beta-activated kinase 1 and MAP3K7-binding protein 1 | 1305 | 54610 | 32 | 32 | 10 | 10 | 1.39 |
| SCYL2 | SCYL2 | SCY1-like protein 2 | 1302 | 103642 | 33 | 33 | 10 | 10 | 0.59 |
| K1C10 | KRT10 | Keratin, type I cytoskeletal 10 | 1286 | 58792 | 42 | 42 | 14 | 14 | 2.37 |
| PIGT | PIGT | GPI transamidase component PIG-T | 1150 | 65658 | 44 | 44 | 15 | 15 | 1.97 |
| KCTD5 | KCTD5 | BTB/POZ domain-containing protein KCTD5 | 1137 | 26076 | 24 | 24 | 5 | 5 | 1.48 |
| K22E | KRT2 | Keratin, type II cytoskeletal 2 epidermal | 978 | 65393 | 27 | 27 | 10 | 10 | 1.08 |
| KCD17 | KCTD17 | BTB/POZ domain-containing protein KCTD17 | 959 | 35648 | 21 | 21 | 6 | 6 | 1.54 |
| GPAA1 | GPAA1 | Glycosylphosphatidylinositol anchor attachment 1 protein | 916 | 67580 | 28 | 28 | 7 | 7 | 0.76 |
| BIP | HSPA5 | Endoplasmic reticulum chaperone BiP | 837 | 72288 | 23 | 23 | 12 | 12 | 1.21 |
| MCM3 | MCM3 | DNA replication licensing factor MCM3 | 817 | 90924 | 30 | 30 | 13 | 13 | 0.98 |
| GPI8 | PIGK | GPI-anchor transamidase | 744 | 45223 | 28 | 28 | 7 | 7 | 1.09 |
| EF1A1 | EEF1A1 | Elongation factor 1-alpha 1 | 721 | 50109 | 24 | 24 | 6 | 6 | 0.77 |
| HSP7C | HSPA8 | Heat shock cognate 71 kDa protein | 656 | 70854 | 21 | 21 | 8 | 8 | 0.72 |
| RS3 | RPS3 | 40S ribosomal protein S3 | 656 | 26671 | 27 | 27 | 8 | 8 | 3.96 |
| EF1D | EEF1D | Elongation factor 1-delta | 643 | 31103 | 13 | 13 | 5 | 5 | 1.15 |
| TTC28 | TTC28 | Tetratricopeptide repeat protein 28 | 640 | 270715 | 19 | 19 | 10 | 10 | 0.19 |
| ACTB | ACTB | Actin, cytoplasmic 1 | 598 | 41710 | 12 | 12 | 5 | 5 | 0.77 |
| M3K7 | MAP3K7 | Mitogen-activated protein kinase kinase kinase 7 | 526 | 67153 | 19 | 19 | 5 | 5 | 0.43 |
| HS71A | HSPA1A | Heat shock 70 kDa protein 1A | 519 | 70009 | 23 | 23 | 9 | 9 | 0.85 |
| EFTU | TUFM | Elongation factor Tu, mitochondrial | 497 | 49510 | 14 | 14 | 5 | 5 | 0.62 |
| GOLP3 | GOLPH3 | Golgi phosphoprotein 3 | 488 | 33790 | 11 | 11 | 6 | 6 | 1.33 |
| MCM5 | MCM5 | DNA replication licensing factor MCM5 | 469 | 82233 | 12 | 12 | 5 | 5 | 0.34 |
| PRP8 | PRPF8 | Pre-mRNA-processing-splicing factor 8 | 460 | 273427 | 25 | 25 | 16 | 16 | 0.32 |
| E2AK3 | EIF2AK3 | Eukaryotic translation initiation factor 2-alpha kinase 3 | 390 | 125137 | 12 | 12 | 6 | 6 | 0.26 |
| SERPH | SERPINH1 | Serpin H1 | 377 | 46411 | 11 | 11 | 5 | 5 | 0.67 |
| KPYM | PKM | Pyruvate kinase PKM | 371 | 57900 | 12 | 12 | 6 | 6 | 0.64 |
| EF1G | EEF1G | Elongation factor 1-gamma | 367 | 50087 | 14 | 14 | 5 | 5 | 0.61 |
| SF3B3 | SF3B3 | Splicing factor 3B subunit 3 | 358 | 135492 | 13 | 13 | 8 | 8 | 0.33 |
| RBBP7 | RBBP7 | Histone-binding protein RBBP7 | 347 | 47790 | 12 | 12 | 6 | 6 | 0.82 |
| RPN1 | RPN1 | Dolichyl-diphosphooligosaccharide--protein glycosyltransferase subunit 1 | 321 | 68527 | 14 | 14 | 7 | 7 | 0.63 |
| PRDX1 | PRDX1 | Peroxiredoxin-1 | 321 | 22096 | 14 | 14 | 6 | 6 | 2.61 |
| RS16 | RPS16 | 40S ribosomal protein S16 | 317 | 16435 | 9 | 9 | 3 | 3 | 1.36 |
| U5S1 | EFTUD2 | 116 kDa U5 small nuclear ribonucleoprotein component | 310 | 109366 | 14 | 14 | 11 | 11 | 0.62 |
| VIME | VIM | Vimentin | 300 | 53619 | 11 | 11 | 5 | 5 | 0.56 |
| RS18 | RPS18 | 40S ribosomal protein S18 | 296 | 17708 | 8 | 8 | 3 | 3 | 1.22 |
| PSA4 | PSMA4 | Proteasome subunit alpha type-4 | 285 | 29465 | 10 | 10 | 3 | 3 | 0.62 |
| JAK1 | JAK1 | Tyrosine-protein kinase JAK1 | 282 | 133191 | 13 | 13 | 9 | 9 | 0.38 |
| STK38 | STK38 | Serine/threonine-protein kinase 38 | 281 | 54155 | 10 | 10 | 5 | 5 | 0.55 |
| OST48 | DDOST | Dolichyl-diphosphooligosaccharide--protein glycosyltransferase 48 kDa subunit | 281 | 50769 | 13 | 13 | 5 | 5 | 0.6 |
| TCPE | CCT5 | T-complex protein 1 subunit epsilon | 277 | 59633 | 8 | 8 | 5 | 5 | 0.49 |
| FLNA | FLNA | Filamin-A | 276 | 280564 | 11 | 11 | 8 | 8 | 0.15 |
| HNRPK | HNRNPK | Heterogeneous nuclear ribonucleoprotein K | 271 | 50944 | 4 | 4 | 2 | 2 | 0.21 |
| EF1B | EEF1B2 | Elongation factor 1-beta | 268 | 24748 | 4 | 4 | 2 | 2 | 0.47 |
| ICLN | CLNS1A | Methylosome subunit pICln | 255 | 26199 | 9 | 9 | 1 | 1 | 0.2 |
| HS90B | HSP90AB1 | Heat shock protein HSP 90-beta | 255 | 83212 | 9 | 9 | 5 | 5 | 0.33 |
| CH60 | HSPD1 | 60 kDa heat shock protein, mitochondrial | 242 | 61016 | 8 | 8 | 6 | 6 | 0.6 |
| TMEDA | TMED10 | Transmembrane emp24 domain-containing protein 10 | 237 | 24960 | 4 | 4 | 3 | 3 | 0.77 |
| RS27A | RPS27A | Ubiquitin-40S ribosomal protein S27a | 230 | 17953 | 6 | 6 | 2 | 2 | 0.69 |
| RS9 | RPS9 | 40S ribosomal protein S9 | 225 | 22578 | 15 | 15 | 6 | 6 | 2.52 |
| OCAD2 | OCIAD2 | OCIA domain-containing protein 2 | 220 | 16943 | 4 | 4 | 2 | 2 | 0.74 |
| KCTD2 | KCTD2 | BTB/POZ domain-containing protein KCTD2 | 217 | 28509 | 6 | 6 | 3 | 3 | 0.65 |
| LRC59 | LRRC59 | Leucine-rich repeat-containing protein 59 | 213 | 34909 | 12 | 12 | 4 | 4 | 0.73 |
| HNRH1 | HNRNPH1 | Heterogeneous nuclear ribonucleoprotein H | 209 | 49198 | 5 | 5 | 3 | 3 | 0.34 |
| RS2 | RPS2 | 40S ribosomal protein S2 | 193 | 31305 | 10 | 10 | 5 | 5 | 1.13 |
| TIF1B | TRIM28 | Transcription intermediary factor 1-beta | 183 | 88493 | 6 | 6 | 3 | 3 | 0.18 |
| TRAP1 | TRAP1 | Heat shock protein 75 kDa, mitochondrial | 176 | 80060 | 5 | 5 | 3 | 3 | 0.2 |
| CREL1 | CRELD1 | Cysteine-rich with EGF-like domain protein 1 | 174 | 45409 | 7 | 7 | 4 | 4 | 0.52 |
| PIGU | PIGU | Phosphatidylinositol glycan anchor biosynthesis class U protein | 168 | 50019 | 2 | 2 | 1 | 1 | 0.1 |
| PRP19 | PRPF19 | Pre-mRNA-processing factor 19 | 167 | 55146 | 6 | 6 | 3 | 3 | 0.3 |
| VAPA | VAPA | Vesicle-associated membrane protein-associated protein A | 165 | 27875 | 4 | 4 | 2 | 2 | 0.41 |
| EF2 | EEF2 | Elongation factor 2 | 159 | 95277 | 6 | 6 | 6 | 6 | 0.35 |
| GRP75 | HSPA9 | Stress-70 protein, mitochondrial | 155 | 73635 | 3 | 3 | 1 | 1 | 0.07 |
| ARHGA | ARHGEF10 | Rho guanine nucleotide exchange factor 10 | 152 | 151516 | 5 | 5 | 4 | 4 | 0.13 |
| MCM7 | MCM7 | DNA replication licensing factor MCM7 | 145 | 81257 | 5 | 5 | 5 | 5 | 0.34 |
| RL18 | RPL18 | 60S ribosomal protein L18 | 145 | 21621 | 2 | 2 | 1 | 1 | 0.24 |
| ASCC3 | ASCC3 | Activating signal cointegrator 1 complex subunit 3 | 139 | 251301 | 6 | 6 | 4 | 4 | 0.08 |
| HAT1 | HAT1 | Histone acetyltransferase type B catalytic subunit | 138 | 49481 | 3 | 3 | 2 | 2 | 0.21 |
| U520 | SNRNP200 | U5 small nuclear ribonucleoprotein 200 kDa helicase | 133 | 244353 | 5 | 5 | 5 | 5 | 0.1 |
| RL7 | RPL7 | 60S ribosomal protein L7 | 126 | 29207 | 6 | 6 | 4 | 4 | 0.92 |
| ANM1 | PRMT1 | Protein arginine N-methyltransferase 1 | 125 | 42434 | 7 | 7 | 4 | 4 | 0.57 |
| RL13 | RPL13 | 60S ribosomal protein L13 | 124 | 24247 | 7 | 7 | 3 | 3 | 0.8 |
| TERA | VCP | Transitional endoplasmic reticulum ATPase | 123 | 89266 | 7 | 7 | 7 | 7 | 0.45 |
| 1433B | YWHAB | 14-3-3 protein beta/alpha | 123 | 28065 | 6 | 6 | 3 | 3 | 0.66 |
| TCPB | CCT2 | T-complex protein 1 subunit beta | 121 | 57452 | 5 | 5 | 3 | 3 | 0.28 |
| RS8 | RPS8 | 40S ribosomal protein S8 | 121 | 24190 | 6 | 6 | 2 | 2 | 0.48 |
| TMED5 | TMED5 | Transmembrane emp24 domain-containing protein 5 | 121 | 25988 | 4 | 4 | 2 | 2 | 0.44 |
| DHX15 | DHX15 | Pre-mRNA-splicing factor ATP-dependent RNA helicase DHX15 | 118 | 90875 | 5 | 5 | 4 | 4 | 0.23 |
| RS6 | RPS6 | 40S ribosomal protein S6 | 118 | 28663 | 4 | 4 | 2 | 2 | 0.39 |
| HNRPU | HNRNPU | Heterogeneous nuclear ribonucleoprotein U | 117 | 90528 | 6 | 6 | 4 | 4 | 0.24 |
| K1C9 | KRT9 | Keratin, type I cytoskeletal 9 | 115 | 62027 | 5 | 5 | 2 | 2 | 0.17 |
| VDAC2 | VDAC2 | Voltage-dependent anion-selective channel protein 2 | 115 | 31547 | 2 | 2 | 1 | 1 | 0.16 |
| ADT2 | SLC25A5 | ADP/ATP translocase 2 | 112 | 32831 | 5 | 5 | 2 | 2 | 0.34 |
| HACD3 | HACD3 | Very-long-chain (3R)-3-hydroxyacyl-CoA dehydratase 3 | 112 | 43132 | 1 | 1 | 1 | 1 | 0.12 |
| SF3B4 | SF3B4 | Splicing factor 3B subunit 4 | 111 | 44357 | 3 | 3 | 2 | 2 | 0.24 |
| TCPH | CCT7 | T-complex protein 1 subunit eta | 111 | 59329 | 2 | 2 | 1 | 1 | 0.08 |
| PRPS1 | PRPS1 | Ribose-phosphate pyrophosphokinase 1 | 109 | 34812 | 3 | 3 | 2 | 2 | 0.31 |
| DDX3X | DDX3X | ATP-dependent RNA helicase DDX3X | 108 | 73198 | 4 | 4 | 4 | 4 |  |
