## Supplementary material for "Uncovering protein-protein interactions of the human sodium channel Nav1.7": Table 2

Table 2. Comparison between mNav1.7- and hNaV1.7-interacted proteins.

| Translational Protein | Homo | **Eef1a1** | Eef1b2 | Eef1d | **Eef2** | Eftud2 | Mrps23 | Mrps34 | Rpl11 | Rpl13 | Rpl14 | Rpl18 | Rpl18a | Rpl23 |
| --- | --- | --- | --- | --- | --- | --- | --- | --- | --- | --- | --- | --- | --- | --- |
|  |  | Rpl27 | Rpl27a | Rpl6 | Rpl7 | Rplp0p6 | Rps11 | Rps15a | Rps16 | Rps17 | Rps18 | Rps2 | Rps25 | Rps3 |
|  |  | Rps3a | Rps4x | Rps5 | Rps6 | Rps8 | Rps9 | **Tufm** |  |  |  |  |  |  |
|  | Mouse | **Eef1a1** | Eef1a2 | **Eef2** | Eif3a | Eif3b | Eif3c | Eif3d | Eif3e | Eif3i | Eif3l | **Tufm** |  |  |
| Chaperone | Homo | Canx | **Cct2** | **Cct3** | **Cct5** | **Cct6a** | **Cct7** | Clgn | Clip1 | **Hsp90ab1** | Hspa1a | **Hspa5** | Hspa8 | **Hspa9** |
|  |  | Npm1 | Pdia6 | **Tcp1** | Trap1 |  |  |  |  |  |  |  |  |  |
|  | Mouse | Ahsa1 | Calr | **Cct2** | **Cct3** | Cct4 | **Cct5** | **Cct6a** | [**Cct7**](http://pantherdb.org/panther/category.do?categoryAcc=PC00073) | Cct8 | Dctn1 | Hsp90aa1 | **Hsp90ab1** | **Hspa5** |
|  |  | **Hspa9** | Hsph1 | Ppia | **Tcp1** |  |  |  |  |  |  |  |  |  |
| Protein Modifying Enzyme | Homo | Akt1 | Carm1 | Eif2ak3 | Hpr | Jak1 | **Map3k7** | **Ogt** | Ppm1b | Prkdc | Prmt1 | Prmt5 | **Psma4** | Scyl2 |
|  |  | Sec11a | **Stk38** | Stub1 | **Tab1** | Trim21 | Trim28 | Ubr5 | Usp7 | Usp9x |  |  |  |  |
|  | Mouse | Ank3 | Asrgl1 | Map2 | **Map3k7** | **Ogt** | Ppm1a | Ppp1cc | Ppp2cb | Psma1 | **Psma4** | Psma5 | Psma7 | Psmb2 |
|  |  | Psmb4 | Psmb5 | Psmc2 | Psmc3 | Psmc4 | Psmc5 | Psmd1 | Psmd12 | Psmd13 | Psmd2 | Psmd3 | Psmd4 | **Stk38** |
|  |  | Stk38l | **Tab1** | Tcp1 | Uchl1 | Uqcrc1 | Uqcrc2 | Usp15 |  |  |  |  |  |  |
| Cytoskeletal Protein | Homo | Actb | Actn1 | **Capza1** | **Cfl1** | Dnah17 | Kif11 | Krt10 | Krt9 | Pls3 | Shroom3 | Sptan1 | Tuba1a | Tuba1b |
|  |  | **Tubb** | Tubb2a | Tubb4a | Tubb4b | Vim |  |  |  |  |  |  |  |  |
|  | Mouse | Ablim1 | Actbl2 | Actr1a | Actr1b | Actr2 | **Capza1** | **Cfl1** | Clasp2 | Cttn | Dbn1 | Dctn4 | Dctn5 | Des |
|  |  | Dsp | Dst | Dstn | Dync1h1 | Dynll2 | Epb41l1 | Epb41l3 | Kif5b | Kif5c | Klc1 | Myo6 | Prph | Tmod2 |
|  |  | Tuba4a | Tubb2b | **Tubb5** |  |  |  |  |  |  |  |  |  |  |
| Rna Metabolism Protein | Homo | Cpsf1 | Cpsf6 | **Ddb1** | Ddx3x | Dhx15 | E2f7 | **Hnrnph1** | Hnrnpk | Larp1 | Med14 | Nop56 | Nudt21 | Pabpc1 |
|  |  | Pcbp1 | Pcf11 | Pdcd11 | Polr2a | Prpf19 | Prpf31 | Prpf6 | Prpf8 | Ptbp1 | Sf3b1 | Sf3b2 | Sf3b3 | Sf3b4 |
|  |  | Sf3b6 | Snrpd2 | Syncrip | Tardbp |  |  |  |  |  |  |  |  |  |
|  | Mouse | **Ddb1** | **Hnrnph1** | Hnrnph2 | Htatsf1 | Sart3 |  |  |  |  |  |  |  |  |
| Intercellular Signal Molecule | Homo | Gdf9 | Tmpo |  |  |  |  |  |  |  |  |  |  |  |
|  | Mouse | Fgb | Fgg |  |  |  |  |  |  |  |  |  |  |  |
| Membrane Traffic Protein | Homo | Clstn3 | Copa | Lman2l | Ociad2 | **Tmed10** | Tmed2 | Tmed5 | Tmed9 | Vapa |  |  |  |  |
|  | Mouse | Ankfy1 | Ap2s1 | Ap3b2 | Bcap31 | Dnm1 | Napa | Napb | Nsf | Stx12 | Syt2 | **Tmed10** |  |  |
| Transfer/Carrier Protein | Homo | Alb | Lrp1 | Slc25a5 |  |  |  |  |  |  |  |  |  |  |
|  | Mouse | Fabp7 | Hba | Tf |  |  |  |  |  |  |  |  |  |  |
| Transmembrane Signal Receptor | Homo | Adgre5 | Or1l8 | Pgrmc1 | Pgrmc2 |  |  |  |  |  |  |  |  |  |
|  | Mouse | Glud1 | Tkt |  |  |  |  |  |  |  |  |  |  |  |
| Transporter | Homo | Abca8 | Abcd3 | Atp5f1a | Atp5mf | Ccar2 | Clns1a | Mtch2 | Pls3 | Praf2 | **Slc25a3** | **Vcp** | **Vdac1** |  |
|  | Mouse | Ap3d1 | Aqp4 | Atp1a1 | Atp1a3 | Atp2a2 | Atp5a1 | Atp5c1 | Atp5d | Atp5i | Atp5j | Atp5j2 | Atp5o | Atp6v1g2 |
|  |  | Kpnb1 | Scn3b | Sfxn3 | Slc12a2 | Slc12a5 | Slc17a7 | Slc1a3 | **Slc25a3** | Slc2a1 | Slc4a1 | Slc4a8 | Slc7a14 | Slc7a5 |
|  |  | Slc8a1 | Usmg5 | **Vcp** | **Vdac1** | Vdac3 |  |  |  |  |  |  |  |  |
| Scaffold/Adaptor Protein | Homo | Ankrd28 | Ascc3 | Ivns1abp | Kctd17 | Kctd2 | Kctd5 | Lrrc59 | Lrrc8e | Snrnp200 | Ttc28 | Wdr87 | **Ywhab** |  |
|  | Mouse | Akap12 | Cyfip2 | Kctd12 | Mapk8ip3 | Mpdz | Mpp2 | Ppfia3 | **Ywhab** | Ywhae | Ywhag | Ywhah | Ywhaq |  |
| Metabolite Interconversion Enzyme | Homo | Acot9 | Acsm6 | Cnp | Cox7a2 | Ddost | Ganab | Hacd3 | Hao1 | Impdh2 | Ldhb | Mt-Co2 | Mthfd1 | Nt5c2 |
|  |  | Pfkfb3 | **Pkm** | **Prdx1** | **Prdx2** | Prdx6 | Prps1 | Prpsap1 | Prpsap2 | Rpn1 | Txn |  |  |  |
|  | Mouse | Acly | Aco2 | Agpat3 | Ampd2 | Cox4i1 | Cox5a | Crmp1 | Dpysl2 | Dpysl3 | Dpysl4 | Dpysl5 | Echs1 | Eno1 |
|  |  | Glud1 | Glul | Gpx4 | Mccc2 | Mthfd1l | Ndufa10 | Ndufa7 | Ndufa9 | Ndufb10 | Ndufb7 | Ndufs7 | Ndufs8 | Ndufv3 |
|  |  | Oxct1 | Pccb | Pfkm | Pgam1 | Pi4ka | **Pkm** | **Prdx1** | **Prdx2** | Sbf1 | Synj1 | Tkt |  |  |
| Calcium-Binding Protein | Homo | Anxa2 | Cpne1 | S100b |  |  |  |  |  |  |  |  |  |  |
|  | Mouse | Calb2 | Calm1 |  |  |  |  |  |  |  |  |  |  |  |
| Defense/Immunity Protein | Homo | C1qbp |  |  |  |  |  |  |  |  |  |  |  |  |
|  | Mouse | Ighg1 | Ighm | Igsf8 | Lsamp | Ntm |  |  |  |  |  |  |  |  |
| Gene-Specific Transcriptional Regulator | Homo | Hdx | Mta1 |  |  |  |  |  |  |  |  |  |  |  |
|  | Mouse | Erh |  |  |  |  |  |  |  |  |  |  |  |  |
| Protein-Binding Activity Modulator | Homo | Arhgef10 | Dock4 | Gnal | Ppp2r1a | Prkag1 | Rab5a | Serpina1 | Serpinb10 | Serpinh1 | Srgap1 |  |  |  |
|  | Mouse | Pebp1 | Rab1a | Rp2 |  |  |  |  |  |  |  |  |  |  |
| Others | Homo | **Fasn** | **Flna** | Krt1 | Krt10 | **Krt2** | Krt71 | Krt9 | **Lmna** | Ppp2r1a | **Tln1** |  |  |  |
|  | Mouse | **Fasn** | **Flna** | **Krt2** | Krt6a | **Lmna** | Ppp1cc | Ppp1r9a | Ppp2cb | **Tln1** | Tln2 |  |  |  |
| **In summary** |  |  |  |  |  |  |  |  |  |  |  |  |  |  |
| Homo | Atp5f1a | Atp5mf |  |  |  |  |  | Canx | **Capza1** | **Tcp1** | **Cct2** | **Cct3** | **Cct5** | **Cct6a** |
| Mouse | Atp5a1 | Atp5c1 | Atp5d | Atp5i | Atp5j | Atp5j2 | Atp5o | Calr | **Capza1** | **Tcp1** | **Cct2** | **Cct3** | Cct4 | **Cct5** |
| Homo | **Cct7** |  |  | Cox7a2 |  | **Ddb1** | **Eef1a1** | Eef1b2 | Eef1d | Eef1g | **Eef2** | **Fasn** | **Flna** | **Hnrnph1** |
| Mouse | **Cct6a** | **Cct7** | Cct8 | Cox4i1 | Cox5a | **Ddb1** | **Eef1a1** | Eef1a2 |  |  | **Eef2** | **Fasn** | **Flna** | **Hnrnph1** |
| Homo | Hnrnpk | Hnrnpm | Hnrnpu | **Hsp90ab1** |  | Hspa1a | **Hspa5** | Hspa8 | **Hspa9** | Kctd2 | Kctd5 | Kif11 |  | Krt1 |
| Mouse | Hnrnph2 |  |  | Hsp90aa1 | **Hsp90ab1** |  | **Hspa5** | **Hspa9** |  | Kctd12 |  | Kif5b | Kif5c |  |
| Homo | Krt10 | **Krt2** | Krt71 | Krt9 | **Lmna** | **Map3k7** | **Pkm** | Ppm1b | Ppp2r1a |  |  | **Prdx1** | **Prdx2** |  |
| Mouse |  | **Krt2** | Krt6a |  | **Lmna** | **Map3k7** | **Pkm** | Ppm1a | Ppp1cc | Ppp1r9a | Ppp2cb | **Prdx1** | **Prdx2** | Psma1 |
| Homo | **Psma4** |  |  | Rab5a | **Slc25a3** | Slc25a5 | **Stk38** |  | **Tab1** | Tab2 | Tab3 | **Tln1** |  | **Tmed10** |
| Mouse | **Psma4** | Psma5 | Psma7 | Rab1A | **Slc25a3** |  | **Stk38** | Stk38l | **Tab1** |  |  | **Tln1** | Tln2 | **Tmed10** |
| Homo | Tmed5 | Tmed9 | Tuba1a | Tuba1b | Tubb2a | Tubb4a | Tubb4b | **Tubb5** | **Tufm** | Usp7 | Usp9x | **Vcp** | **Vdac1** | Vdac2 |
| Mouse |  |  | Tuba4a |  | Tubb2b |  |  | **Tubb5** | **Tufm** | Usp15 |  | **Vcp** | **Vdac1** | Vdac3 |
| Homo | **Ywhab** |  |  |  |  |  |  |  |  |  |  |  |  |  |
| Mouse | **Ywhab** | Ywhae | Ywhag | Ywhah | Ywhaq |  |  |  |  |  |  |  |  |  |
